# Wild songbirds exhibit consistent individual differences in inter-specific social behaviour

**DOI:** 10.1101/746545

**Authors:** F. Hillemann, E. F. Cole, D. R. Farine, B. C. Sheldon

## Abstract

1. Natural populations and communities consist of individuals that differ in their phenotypes. There is increasing evidence in community ecology that consistent intraspecific variation in behaviour changes the outcome of ecological interactions.
2. Differences in intra- and inter-specific interactions are expected to play a major role in determining patterns of species coexistence and community structure. However, the question of whether individuals vary in their propensity to associate with heterospecifics has been neglected.
3. We used social network analysis to characterise pattern of heterospecific associations in wild mixed-species flocks of songbirds, and assessed whether individuals adopt consistent social strategies in their broader, heterospecific, social environment. We quantified heterospecific foraging associations using data from a large automatically monitored PIT-tagged population of birds, involving more than 300 000 observations of flock membership, collected over three winters, for two tit species (Paridae), blue tits, *Cyanistes caeruleus*, and great tits, *Parus major*.
4. We assessed individual consistency in interspecific social preferences over both short-term (week-to-week) and longer-term (year-to-year) timescales for a total of 4610 individuals, and found that blue tits and great tits exhibited marked and consistent intraspecific differences in heterospecific social phenotypes in terms of both absolute and relative number of associates. Further, we found that these consistent differences were significantly greater than expected from spatial and temporal differences in population densities.
5. Heterospecific associations represent a major component of the social environment for many species, and our results show that individuals vary consistently in their social decisions with respect to heterospecifics. These findings provide support for the notion that intraspecific trait variation contributes to patterns at community and ecosystem levels.

## INTRODUCTION

The social environment is an important component of individuals’ lives. How individuals interact with conspecifics has been shown to have implications for their susceptibility to disease (Silk et al. 2018), access to resources (Aplin et al. 2012), and reproductive success and survival (Alberts 2019). Many potential mechanisms might mediate a sociality-fitness link, including access to resources, agonistic support, and anti-predator benefits (Ostner & Schülke 2018). While much of our current understanding of the costs and benefits of group living results from research on single-species groups (Krause & Ruxton 2002), social groups can take many different forms, and frequently include individuals from different species (Goodale et al. 2017). Mixed-species groups are widespread across animal communities (amphibians: Glos et al. 2007, birds: Sridhar et al. 2009, fish: Lukoschek & McCormick 2000, mammals: Stensland et al. 2003), and offer opportunities for studying the causes of social behaviour without confounding effects of kin-selection or mate-choice (Farine et al. 2012). Many of the key processes thought important for group-living in single-species contexts have been suggested to also apply to the formation of mixed-species groups (Dhondt 2012, Sridhar & Guttal 2018). For example, both grouping and use of heterospecific social information have been shown to provide anti-predator benefits (Magrath et al. 2015; Meise et al. 2018; Goodale et al. 2019) and foraging benefits (Dolby & Grubb 1998; Farine et al. 2015) to members of mixed-species groups. In turn, interactions with heterospecifics could influence the positions of individuals within their social networks, and thereby contribute to ecological and evolutionary processes arising via the social environment (see Cantor et al. 2019 for a review).

Ecological research on species coexistence has traditionally focussed on averaged differences between species, thus ignoring the importance of intraspecific variation among individuals for species interactions (Bolnick et al. 2011; Violle et al. 2012; Hart et al. 2016). It has become increasingly apparent that individual-level processes are of vital importance for understanding community dynamics and other emerging properties of individual interactions. To study social behaviour in mixed-species groups, Farine et al. (2012) suggested a paradigm shift, from a species-level perspective to treating interactions between individuals as the basic unit of analysis (*sensu* Hinde 1976). Such a bottom-up approach accounts for the effect of inter-individual differences in traits and has long been applied to study sociality in single-species groups, thus providing a theoretical and analytical basis for analysing heterospecific sociality. Social Network Analysis (SNA) has become an important method for quantifying individuals’ social decisions (such as partner choice), interaction pattern among individuals, and the emerging social structure of groups and populations (Cantor et al. 2019). To our knowledge, only two studies have analysed dyadic interactions to explore the role of individual-level decisions in shaping heterospecific groups and both focus on within-species variation in measures of social behaviour (Farine et al. 2012; Farine & Milburn 2013). These studies show that a bottom-up approach is useful when exploring hypotheses regarding individual variation in heterospecific association propensity. Yet, neither of these studies explored whether individuals differ consistently in their propensity to associate with heterospecifics.

Considering individuals’ social decisions in a broader, multi-species context will allow a better understanding of the link between individual-level decisions, fitness-related outcomes, and the structure of mixed-species social communities (Farine et al. 2012). For example, how individuals are positioned within their social environment can predict their reproductive success (Formica et al. 2011; Farine & Sheldon 2015). Specifically, the propensity for great tits, *Parus major*, to acquire a breeding territory has been shown to be higher, not only for birds that dispersed into the population earlier, but also when individuals dispersed early relative to their competitors (Farine & Sheldon 2015). For selection to act via the social environment, individuals must vary consistently in their social environment (McDonald et al. 2017). Hence, a key first step in relating fitness consequences to social network position is to understand the distribution and consistency of individual variation in behaviour by characterising individual interaction patterns and social phenotypes. Consistency in social behaviour is often assumed, but only a few studies have shown that individuals can express consistent social phenotypes, by repeatedly measuring and comparing their mebehaviour over larger time spans (Blumstein et al. 2013; Jacoby et al. 2014; Aplin et al. 2015; Menz et al. 2017; O’Brien et al. 2018). Whether individuals also express consistent social strategies in a multi-species context is yet to be explored.

In this study, we test whether individual members of wintering mixed-species flocks of songbirds have consistent interspecific social preferences. Over three winters, we recorded associations among two tit species (Paridae) fitted with individual passive integrated transponder (PIT) tags, recorded by a grid of automated feeding stations fitted with radio frequency identification (RFID) antennae. This large-scale observational study provided a unique opportunity to assess individual consistency in interspecific social preferences over both short-term (week-to-week) and longer-term (year-to-year) timescales. We used SNA to quantify measures of gregariousness and heterospecific flocking propensity for 2775 individual blue tits, *Cyanistes caeruleus*, and 1835 great tits, observed over three years, and then assessed the individual consistency of these traits using repeatability analyses. We then used network randomisation approaches (Farine 2017) to calculate repeatability estimates after accounting for spatio-temporal differences in the distribution, and thus broad social environments, among individuals. By comparing the two sets of repeatability estimates, we were able to separate consistency in social decision making from consistency in behavior caused by spatial factors.

## METHODS

### Study system

This study was conducted from December 2011 to March 2014, in the context of a long-term research project studying the social behaviour of tits in Wytham Woods, Oxfordshire, U.K. (51°46’N, 01°20’W). Blue tits and great tits, as well as some coal tits, *Parus ater*, marsh tits, *Poecile palustris*, and European nuthatches, *Sitta europaea*, were individually tagged with a unique British Trust for Ornithology (BTO) metal leg band, and a plastic leg ring carrying a passive integrated transponder (PIT tag, IB Technologies, UK). Birds were ringed as nestlings or as adults caught at nest-boxes during the breeding season, or using mist-nets in winter. Sex and age (first year/adult) were identified using plumage coloration or breeding records. The proportion of the population being tagged was very high, estimated over 90% for the first year of the study (see Aplin et al. 2013 for a formal analysis).

Tits form mixed-species foraging flocks with fission-fusion dynamics outside the breeding season, which mainly consist of Parid species, but can be joined by nuthatches, treecreepers, and woodpeckers (Hinde 1952; Ekman 1989). In this study, we consider heterospecific interactions between blue tits and great tits, the most abundant species in this population, that together account for 90.6 ± 0.8 % (mean ± SEM) of individuals recorded across the three winters under study.

### Data collection

We collected data on social behaviour of individual birds by recording their visits to feeding stations equipped with radio-frequency identification (RFID) antennae (Dorset ID, Netherlands). We deployed a total of 65 sunflower-seed bird feeders with two access points were deployed throughout the study area in an even grid of approximately 250 × 250 m (see Supplementary Material Figure S1). These feeding stations opened automatically before sunrise and shut down after dusk on Saturday and Sunday (hereafter *weekends*), and remained closed on weekdays. Each time a bird visited the feeder, the identity of the bird detected by the RFID antenna would be logged automatically onto the RFID logger, and we downloaded these data from each feeder weekly. Data for this study was collected during three separate seasons of 14 weekends each, over three winters: 3 December 2011 to 27 February 2012 (Year 1), 1 December 2012 to 3 March 2013 (Year 2), and 30 November 2013 to 2 March 2014 (Year 3). These time periods correspond to those used to assess individual differences social phenotypes of great tits in a single-species context (by Aplin et al. 2015). Number of individuals observed in each season is given in Table 1.

**Table 1.** Number of individual great tits and blue tits included in each winter data collection period, and spread of metrics of social behaviour. Annual mean values across all individuals are provided, and 1^st^ and 3^rd^ quartile in brackets.

|  |  | Blue tit | Great tit |
| --- | --- | --- | --- |
| **N individuals** | 2011-12 | 1631 | 1085 |
|  | 2012-13 | 1306 | 813 |
|  | 2013-14 | 1154 | 842 |
| **Degree** | 2011-12 | 131.2 [71.0, 183.0] | 169.9 [106.0, 226.0] |
|  | 2012-13 | 101.9 [49.0, 145.0] | 115.1 [65.0, 158.0] |
|  | 2013-14 | 93.6 [50.5, 133.0] | 111.3 [62.0, 156.0] |
| **Strength** | 2011-12 | 3.8 [1.2, 5.8] | 5.0 [3.4, 6.8] |
|  | 2012-13 | 3.2 [0.9, 5.1] | 4.2 [1.9, 6.2] |
|  | 2013-14 | 2.9 [1.1, 4.4] | 3.4 [1.7, 5.0] |
| **Group size** | 2011-12 | 11.6 [9.0, 14.0] | 11.6 [9.5, 13.8] |
|  | 2012-13 | 10.3 [6.9, 13.2] | 9.9 [6.9, 13.0] |
|  | 2013-14 | 8.8 [6.1, 11.1] | 8.7 [6.2, 10.9] |
| **E-I index** | 2011-12 | -0.05 [-0.18, 0.68] | -0.06 [-0.20, 0.07] |
|  | 2012-13 | -0.10 [-0.24, 0.04] | 0.02 [-0.10, 0.13] |
|  | 2013-14 | -0.06 [-0.19, 0.08] | -0.05 [-0.19, 0.10] |

### Detecting groups

We followed the same protocol as previous studies on this system (e.g., Aplin et al. 2013, 2015; Farine et al. 2012, 2015) to process the datastream of spatiotemporal detections of PIT-tagged birds at feeders. We used a Gaussian mixture model (GMM) to identify gathering events, or flocks (Psorakis et al. 2012). The GMM works by identifying short-term bursts of activity of individuals repeatedly visiting a feeder. When birds visit a feeder, they briefly perch to collect a seed that they then take to nearby vegetation to process (see Supplementary Material Video S1). These visits generate temporal waves in the number of detections that the GMM identifies. The advantage of the GMM approach is that it can identify flocks of differing sizes, and modelling shows that the resulting networks are more robust than other approaches (Psorakis et al. 2015). We applied this method to the data at each feeder on each day, and aggregated data from all of the flocks into one group by individual matrix for each of the 42 weekends in our study period. Detection of groups was done using the gmmevents function in the asnipe package (Farine 2013) in R v.3.4.4 (R Core Team, 2012)

### Constructing social networks

Dyadic association strength was calculated using the simple ratio index. The simple ratio index generates a value between 0 and 1, which represents to probability of observing two individuals together given that at least one individual was in a given flock (Hoppitt & Farine 2017). This represents an unbiased estimate of the proportion of times two individuals spend together (Whitehead 1995). We combined association strengths for each dyad for each weekend to produce a social network. We generated 42 weekend social networks. In addition, we combined data on flock membership over whole winters to produce on social network for each season (3 winter). Social networks were constructed using the asnipe package (Farine 2013) in R v.3.4.4 (R Core Team, 2012).

### Measures of social phenotype

We calculated the following individual-level measures of social behaviour for great tits and blue tits: i) measures of gregariousness: unweighted degree (total number of associates of any species), weighted degree (overall association strength for all species interactions), and average foraging group size, and ii) a relative measure of heterospecific flocking propensity that is independent from an individual’s gregariousness: a variant of the ‘external-internal’ (E-I) index (Krackhardt & Stern 1988). The E-I index quantifies the relationship between links in two exclusive categories and was originally proposed for friendship links to organisational subunits (i.e., external: friendship links between subunits, internal: friendship links within subunits). Analogously, we calculated the E-I index by subtracting the number of ties to conspecifics from the number of heterospecific associations, and dividing this by the total number of associations. Values range from 1 to −1, with 1 specifying that the focal only associates with heterospecifics, −1 indicating strong homophily (focal individual only has ties to conspecifics); the index will be zero if links are distributed equally between conspecifics and heterospecifics.

### Repeatability Analysis

We calculated the proportion of phenotypic trait variation that can be attributed to inter-individual variation, where repeated samples corresponded to replicated networks. To assess consistency in social phenotypes over short and long time periods, we calculated within-year (week-to-week) and between-year repeatability in measures of great tit and blue tit behaviour. Our approach to analysing repeatability of heterospecific social behaviour follows methods previously used to calculate individual differences in the social phenotypes of great tits in a single-species context (Aplin et al. 2015). We calculated repeatability scores using linear mixed-effects models, as the proportion of variance that can be explained by the individual random effect (intra-class correlation coefficient, ICC; Nakagawa & Schielzeth 2010). Our models accounted for the global population size and network density of the sampling period (weekend or season, respectively). For all models, with the exception of the E-I index, we square-rooted the response variables to conform to normality. We estimated a 95% range confidence intervals of the repeatability scores using restricted maximum likelihood Markov chain Monte Carlo sampling, using the R package MCMCglmm with default priors (Hadfield 2010).

### Comparing observed repeatability results to null expectations

As is often the case for social networks, we can expect strong influences of i) resource distribution and other spatial features on group formation (Farine & Sheldon 2019), and ii) local population density (Farine et al. 2015), on social metrics. We therefore used network randomisation approaches (Farine 2017) to control for the spatio-temporal distribution of individuals to isolate the social component of individuals’ decisions from environmental factors. We designed a null model that randomised identities in each step (1000 randomisations in total), but controlled for the spatio-temporal distribution of individuals and species identity. Within each network, we randomly swapped identities among individuals (node permutations), but restricted swaps to be between individuals of the same species that had the majority of their visits recorded at the same feeding station during the data collection for that network. Such restricted node permutations were necessary in order to maintain the total amount of variance in the model constant when applied to randomised datasets by maintaining a consistent social network structure, while randomising the link within individuals across different networks. We estimated the repeatability values for each of the 1000 resulting randomised networks using the same mixed-model approach described above. From the distribution of null repeatability values, we extracted the 95% to represent the expected range of individual behaviour without social preferences, i.e. the repeatability arising from their spatiotemporal occurrence only. To calculate what proportion of the repeatability was accounted for by the permuted data, we divided the mean of the randomised estimates by the observed repeatability estimate.

### Ethical Note

This work was part of an ongoing long-term research project at Wytham Woods, which was approved by the local ethical review panel at the Department of Zoology, University of Oxford. All catching, handling, and ringing of birds was conducted by experienced BTO licence holders.

## RESULTS

We quantified heterospecific associations in 342 510 observations of flock membership, or grouping events, recorded over three years. Great tits were on average observed in 13.57 ± 10.26 (mean ± SD) sampling periods (weekends), and blue tits in 11.37 ± 10.03 sampling periods, across the three years. The majority of birds were recorded in one winter only (1876 blue tits and 1270 great tits), but 629 blue tits and 356 great tits were observed in two years, and 270 blue tits and 209 great tits were recorded in all three years of data collection.

Blue tits and great tits exhibited considerable intraspecific variation in measures of social behaviour (see Table1). On average, individuals had an annual total of 120 associates, but some individuals were recorded foraging with more than 600 other individuals in one year. Birds also varied markedly in the average strength to their associations, with some individuals being three-times more strongly connected to their associates than the population’s mean edge strength. Blue tits and great tits were on average seen with ten other group members, but some individuals’ annual mean foraging group sizes reached up to 40. The relative number of heterospecific to conspecific group members (E-I index) also varied considerably for individuals, with some preferentially associating with conspecifics (negative values of E-I index) and others preferentially associating with heterospecifics (positive values of E-I index). Table 1 summarises the spread of inter-individual differences in measures of social behaviour, and provides number of individuals included in each year.

Individuals from both species were significantly repeatable in their heterospecific social behaviour on both short (week-to-week) and long (year-to-year) timescales (Figure 1, Table S1). Across the three years of the study, the average week-to-week repeatability scores ranged from 0.49 to 0.62 for heterospecific degree, from 0.43 to 0.61 for heterospecific edge strength, from 0.45 to 0.63 for heterospecific average foraging group size, and from 0.38 to 0.51 for E-I index. Table S1 in the Supplementary Material lists repeatability scores separately for blue tits and great tits. Year-to-year repeatability estimates were similar to, or slightly higher than within-year scores for degree, edge strength, and group size, but the between-year consistency in relative number of conspecific and heterospecific associates (E-I index) was moderately low for both blue tits and great tits, with R=0.38 and R=0.25, respectively.

**Fig. 1:**
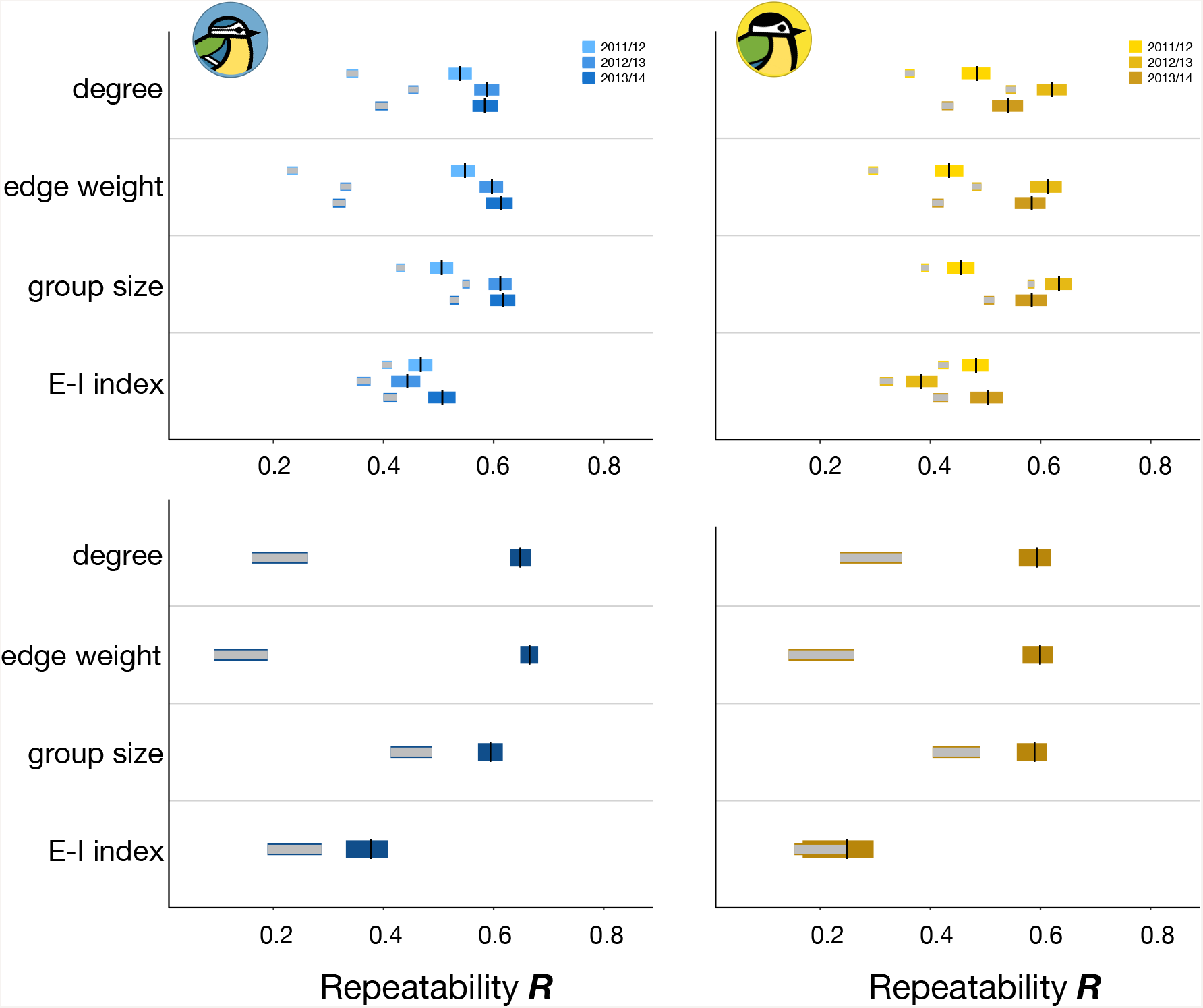
Repeatability scores for week-to-week (top row) and between-year (bottom row) heterospecific social behaviour in great tits (left) and blue tits (right). Black lines show the R score estimates, and yellow or blue bars indicating the 95% range, which was obtained using restricted maximum likelihood Markov chain Monte Carlo sampling. Grey bars represent the 95% range of the repeatability values calculated from 1000 spatially-controlled data randomisations. All measures of social behaviour were significantly repeatable (confidence intervals do not overlap with zero; p<0.001 in all cases), and observed values are significantly higher than the repeatability scores from the randomised data (p=0.027 for E-I index between-year repeatability in great tits, and p<0.001 elsewhere). Underlying data can be found in Table S1 in the Supplementary Material.

All repeatability scores were significantly higher than expected by chance. The observed repeatability values were outside the 95% range of estimates calculated from 1000 permuted datasets for each measure, despite the spatially constrained null model sometimes explaining a large share of individuals’ repeatability scores (see Table S1 in the Supplementary Material). For example, where individuals foraged explained 85-90% of blue tits’ week-to-week consistency in average group size, whereas the permuted data only accounted for about 50% of the week-to-week repeatability in blue tits’ association strength. In both blue tits and great tits, the proportion of the individual week-to-week repeatability that can be explained by the spatio-temporal distribution of birds was largest for group size, slightly lower for E-I index, and smallest for degree and edge strength. In other words, how individuals are distributed in space and time contributed relatively more to the consistency in measures of social behaviour for some phenotypes than for others. The repeatability estimates for degree, edge weight, and group size were usually higher when considering year-to-year repeatability, compared to week-to-week repeatability. In contrast, the year-to-year repeatability of individuals’ relative number of conspecific to heterospecific group members was lower than its short-term consistency, from week-to-week.

## DISCUSSION

We show that individual great tits and blue tits differ consistently in their propensity to associate with heterospecifics. In part, the repeatability in heterospecific associations was driven by spatial variation in population densities and grouping tendencies, with 40-90% of the within-year variation and 20-80% of the between-year repeatability in individual behaviour explained by the spatio-temporal distribution of individuals. These high values suggest that individuals may be making decisions about heterospecific associations as a parameter of their habitat choice. Further, we identified a significant social component to the consistency in measures of heterospecific social behaviour over and above the variation explained by space alone, suggesting that individuals make choices about who to associate with on a flock-by-flock basis, or over timescales of minutes. Our results demonstrate that social phenotypes translate into the heterospecific social environment, with individuals showing marked inter-individual variation in overall gregariousness, and connectedness to heterospecifics in multi-species networks. Given the growing evidence for the effects of individuals’ social environment on modulating evolutionary and ecological processes (Cantor et al. 2019), the importance of decisions about the heterospecific social associations could be widely under-appreciated.

While blue tits and great tits were consistent in measures of their heterospecific social behaviour over both short (weekend-to-weekend) and long (year-to-year) timescales, repeatability scores for degree, edge weight, and group size tended to be higher when measured across years, compared to within years. Similarly, between-year repeatability estimates of great tit social phenotypes in a single-species context were higher than their within-year estimates (Aplin et al. 2015). Annual averages of behavioural measures might provide a more accurate description of individual phenotypes than parameters extracted from weekend-long social networks. It is also possible that more consistent individuals have a higher probability of year-to-year survival. These hypotheses are not mutually exclusive and require further investigation.

Direct comparisons of repeatability values, for example between great tits and blue tits, or between different years, are not straightforward, because the calculation of between-individual variance will depend on population sizes of both species, and therefore opportunities to associate with conspecifics and heterospecifics. Yet, compared to repeatability of different behaviours observed in other studies (reviewed in Bell et al. 2009), blue tits and great tits showed a high consistency in their heterospecific social behaviour. For example, repeatability scores reported for migration, mate preference, and parental behaviours were on average less than 0.3, whereas week-to-week repeatability values observed in this study ranged from 0.4 to 0.6, with. Consistent individual differences in social network position were previously shown in only a few populations where social behaviour was studied in a conspecific context (e.g., Jacoby et al. 2014; Menz et al. 2017; O’Brien et al. 2018), including a study on our system, which noted inter-individual differences and reported similar repeatability in great tit social strategies (group size: R = 0.43-0.64, degree: R = 0.46-0.61, association strength: R = 0.41-0.64) in a single-species context (Aplin et al. 2015). Here, we extend findings of studies on single-species systems by showing that individuals vary in their heterospecific social associations in much the same way as was previously reported within species.

A major ecological and evolutionary question is how social groups form. While we found that individual-level repeatability was significantly larger than expected by chance, the null distribution of repeatability values (from the spatially-constrained permuted datasets) were also consistently larger than 0, and within the range of what are considered moderate to high repeatability values for behaviour (Bell et al. 2009). Different metrics of individuals’ heterospecific social phenotypes can thus be explained by differences in spatial choices and within-location social choices to varying relative amounts. Our results suggest that social choices explain a relatively larger proportion of measures of sociability that represent the outcome of repeated measures of associations among the same individuals (e.g., average association strength), while the distribution of individuals, or their choice of location, explains relatively more of the measures that represent cumulative observations (e.g., E-I index). The high levels of repeatability in measures represented by cumulative observations suggest that individuals may be choosing these environments via larger-scale social decisions, such as those involving the decision of where to disperse. Studies on dispersal have found that the presence (Doligez et al. 2003) and behaviour (Seppänen & Forsman 2007) of heterospecifics can play a major role in shaping individual-level dispersal and habitat choice decisions. Thus, the social environment that individuals experience depends on social choices made at different temporal and spatial scales.

Our findings open up many new questions regarding the evolutionary ecology of behaviour in natural populations. Heterospecific associations might have long-term impacts on individual fitness-related traits, going beyond the immediate drivers arising from foraging and anti-predator benefits. For example, a recent analysis on our study system suggest that great tit breeding performance is affected by their heterospecific breeding neighbourhoods (Roth 2019). How differences in individual association pattern can shape the local structure of populations is demonstrated by carry-over effects of social behaviour. For example, associations among wild zebra finches, *Taeniopygia guttata* during the nesting period predict pattern of interactions in subsequent years (Brandl et al. 2019), while winter social networks among great tits translate to breeding neighbourhoods during spring (Firth & Sheldon 2016). The patterns of local structure arising from interactions among individuals can then translate into local communities, and, in turn, these communities can generate repeatable hierarchical social structures (Farine & Sheldon 2019). Thus, carry-over effects of social structure arising from heterospecific social associations warrants much greater attention.

Our study contributes to the emerging picture that a bottom-up approach integrating the different components, or scales, of social decision-making is needed to gain full understanding of how social communities are formed and maintained. Individuals are not only considering conspecifics when making decisions about which groups to join, but also heterospecifics. This insight opens opportunities to address new questions, such as whether non-random social associations form between individuals of different species, how these emerge and are maintained, and their consequences across different ecological and evolutionary processes. Future work should also explore the underlying causes and fitness consequences of individual differences in the propensity to associate with heterospecifics, and test how such processes can influence the evolution of social groups.

## ACKNOWLEDGEMENTS

We are grateful to all members of the EGI’s (Edward Grey Institute of Field Ornithology) Wytham Tit Project and Social Networks Group for their help with the data collection in the field. The work was supported by grants from the European Research Council (AdG 250164) and the BBSRC (BB/L006081/1) awarded to BCS, DRF was funded by the Max Planck Society and the DFG Centre of Excellence 2117 “Centre for the Advanced Study of Collective Behaviour” (ID: 422037984), and FH was funded by a NERC studentship award, ref. 1654580. Many thanks to Raul Costa-Pereira for much appreciated input, and to Josh Firth for help with the analysis and valuable discussion throughout the preparation of the manuscript.

## AUTHOR CONTRIBUTIONS STATEMENT

All authors conceived the idea of the study, FH and DRF analysed the data; FH led the writing of the manuscript. All authors contributed critically to the drafts and gave final approval for publication.

## DATA AVAILABILITY STATEMENT

Data and code used for analyses will be available from the figshare repository.

## SUPPLEMENTARY MATERIAL

**Table S1:** Repeatability scores (R_obs_) and 95% range of confidence, as per restricted maximum likelihood Markov chain Monte Carlo sampling, for degree (total number of associations), edge weight (association strength), average group size, and E-I index. All measures are significantly repeatable and observed values are significantly higher than expected repeatability scores (R_ran_) from 1000 permutations of the data (p=0.027 for E-I index between-year repeatability in great tits, and p<0.001 elsewhere). The spatially constrained null model accounts for different proportions of the within-individual consistency in heterospecific social behaviour.

| Species | Metric | Season | $R_{obs}$ [$R_{obs,low}$ - $R_{obs,up}$] | [$R_{ran,low}$ - $R_{ran,up}$] | $R_{ran,mean}/R_{obs}$ |
| --- | --- | --- | --- | --- | --- |
| Blue tit | Degree | 2011-12 | 0.54 [0.52-0.56] | [0.33-0.35] | 63.66 % |
|  |  | 2012-13 | 0.59 [0.56-0.61] | [0.44-0.46] | 77.20 % |
|  |  | 2013-14 | 0.58 [0.56-0.61] | [0.38-0.41] | 67.83 % |
|  |  | year-to-year | 0.65 [0.63-0.67] | [0.16-0.27] | 32.67 % |
|  | Edge weight | 2011-12 | 0.55 [0.52-0.57] | [0.22-0.24] | 42.79 % |
|  |  | 2012-13 | 0.60 [0.57-0.62] | [0.32-0.34] | 55.57 % |
|  |  | 2013-14 | 0.61 [0.59-0.63] | [0.31-0.33] | 52.20 % |
|  |  | year-to-year | 0.67 [0.65-0.68] | [0.09-0.19] | 21.14 % |
|  | Group size | 2011-12 | 0.51 [0.48-0.53] | [0.42-0.44] | 85.27 % |
|  |  | 2012-13 | 0.61 [0.59-0.63] | [0.54-0.56] | 89.91 % |
|  |  | 2013-14 | 0.62 [0.59-0.64] | [0.52-0.54] | 85.62 % |
|  |  | year-to-year | 0.59 [0.57-0.61] | [0.41-0.49] | 75.86 % |
|  | E-I index | 2011-12 | 0.47 [0.44-0.49] | [0.40-0.42] | 87.05 % |
|  |  | 2012-13 | 0.44 [0.41-0.47] | [0.35-0.38] | 82.10 % |
|  |  | 2013-14 | 0.51 [0.48-0.53] | [0.40-0.42] | 81.38 % |
|  |  | year-to-year | 0.38 [0.33-0.41] | [0.19-0.29] | 63.20 % |
| Great tit | Degree | 2011-12 | 0.49 [0.46-0.51] | [0.35-0.37] | 74.78 % |
|  |  | 2012-13 | 0.62 [0.59-0.65] | [0.54-0.55] | 88.11 % |
|  |  | 2013-14 | 0.54 [0.51-0.57] | [0.42-0.44] | 79.76 % |
|  |  | year-to-year | 0.59 [0.56-0.62] | [0.23-0.34] | 49.23 % |
|  | Edge weight | 2011-12 | 0.43 [0.41-0.46] | [0.29-0.30] | 68.27 % |
|  |  | 2012-13 | 0.61 [0.58-0.64] | [0.48-0.49] | 78.98 % |
|  |  | 2013-14 | 0.58 [0.55-0.61] | [0.40-0.42] | 70.87 % |
|  |  | year-to-year | 0.60 [0.57-0.62] | [0.14-0.26] | 33.61 % |
|  | Group size | 2011-12 | 0.45 [0.43-0.48] | [0.38-0.40] | 85.86 % |
|  |  | 2012-13 | 0.63 [0.61-0.66] | [0.58-0.59] | 92.06 % |
|  |  | 2013-14 | 0.58 [0.55-0.61] | [0.50-0.52] | 86.74 % |
|  |  | year-to-year | 0.59 [0.56-0.61] | [0.40-0.49] | 75.85 % |
|  | E-I index | 2011-12 | 0.48 [0.46-0.51] | [0.41-0.43] | 87.73 % |
|  |  | 2012-13 | 0.38 [0.36-0.41] | [0.31-0.33] | 83.89 % |
|  |  | 2013-14 | 0.50 [0.47-0.53] | [0.41-0.43] | 83.05 % |
|  |  | year-to-year | 0.25 [0.19-0.30] | [0.15-0.25] | 80.94 % |

**Figure S1:**
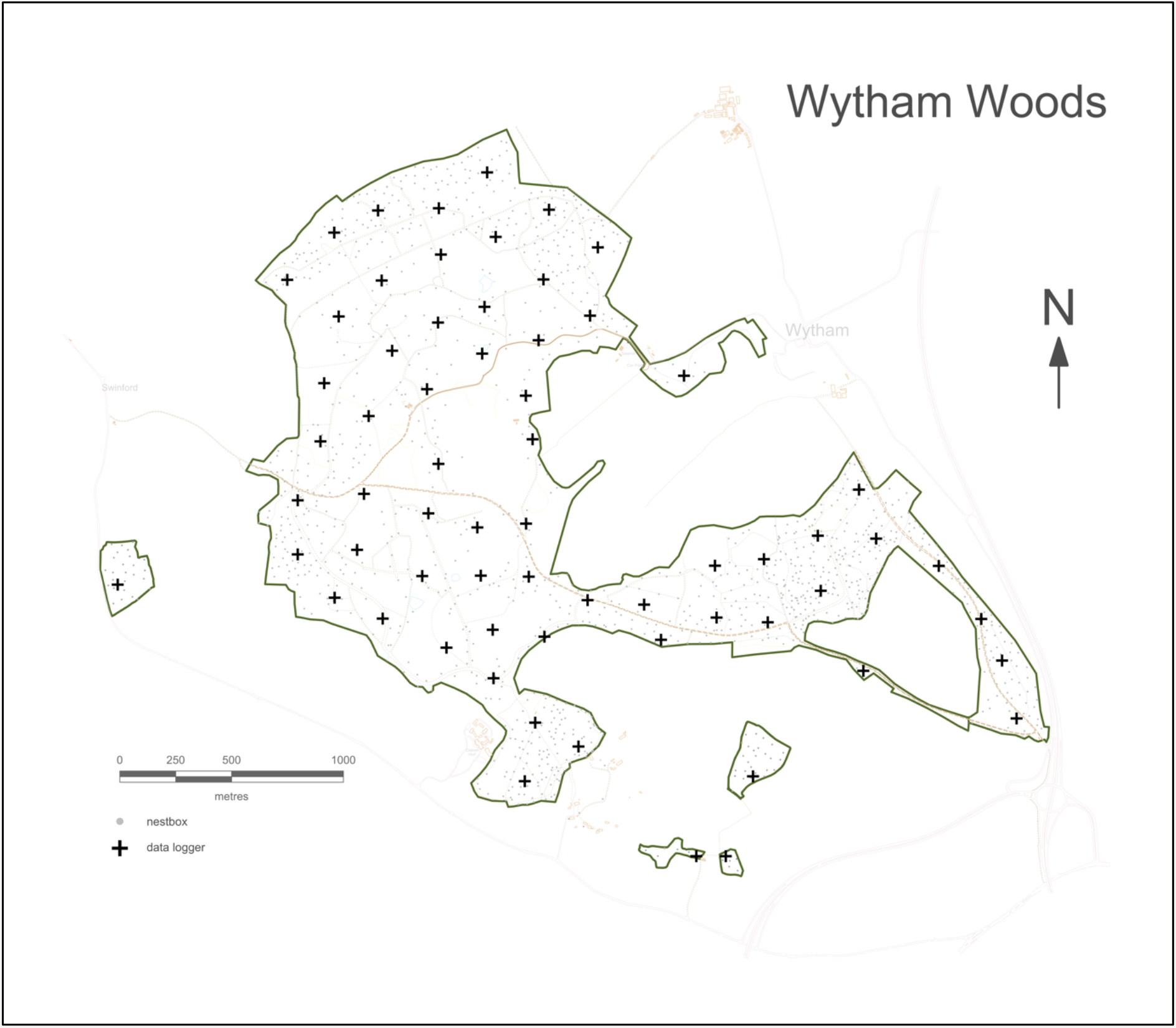
Map of the study site, Wytham Woods (Oxfordshire, UK). Black crosses show the location of the 65 automated RFID feeding stations, which record feeder visits of PIT-tagged birds. These data loggers are distributed in a grid of each approximately 250 × 250 m, and open and close simultaneously on two days per week during the winter months, thus providing a snapshot of the spatio-temporal distribution of birds.

**Video V1:**
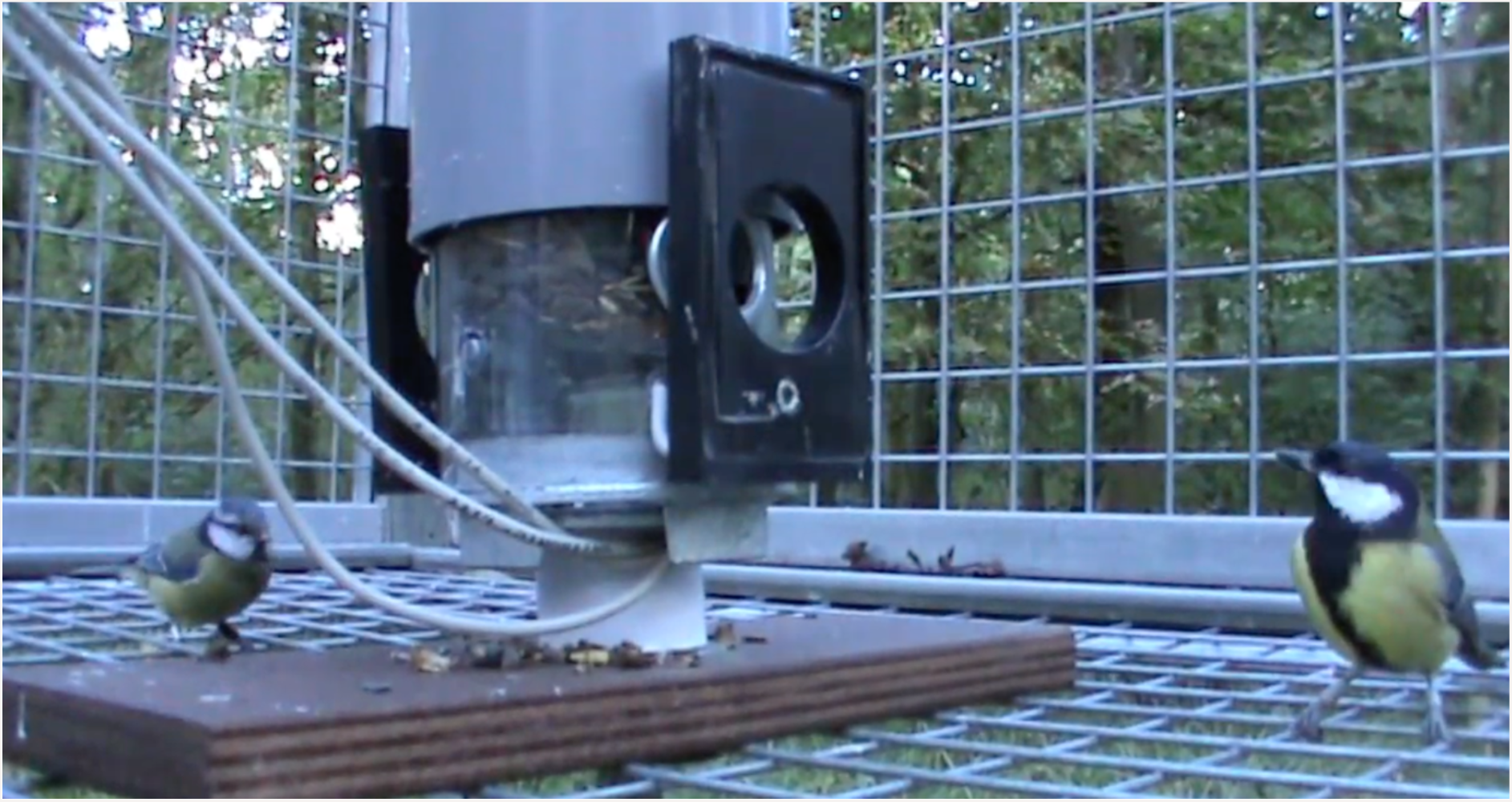
Link to Video demonstrating the collection of social association data at automated feeding station. Birds (Paridae spp.) with PIT-tags moulded into plastic rings on their legs perch on the RFID antennae when taking a seed from one of the feeder’s two access holes, and typically process the unhusked seeds in the nearby vegetation.

